# Insights into the susceptibility of rice to a floral disease

**DOI:** 10.1101/2021.03.03.433744

**Authors:** Guo-Bang Li, Jing Fan, Jie Liu, Jin-Long Wu, Xiao-Hong Hu, Jia-Xue He, Shuai Shen, He Wang, Yong Zhu, Feng He, Han Gao, Zeeshan Ghulam Nabi Gishkori, Jing-Hao Zhao, Yan Li, Fu Huang, Yan-Yan Huang, Zhi-Xue Zhao, Ji-Wei Zhang, Shi-Xin Zhou, Mei Pu, Xuewei Chen, Jing Wang, Weitao Li, Xian-Jun Wu, Yuese Ning, Wenxian Sun, Wen-Ming Wang

## Abstract

Crop floral diseases are economically important as they reduce grain yield and quality and even introduce food toxins. Rice false smut has emerged as a serious floral disease producing mycotoxins. However, very little is known on the interaction mechanisms between rice flower and the causal fungus *Ustilaginoidea virens*. Here we show that a conserved anti-fungal immunity in rice flower is disarmed by *U. virens* via a secreted protein UvChi1. UvChi1 functioned as an essential virulence factor and directly interacted with the chitin receptor CEBiP and co-receptor CERK1 in rice to disrupt their oligomerizations and subsequent immune responses. Moreover, intraspecific-conserved UvChi1 could target OsCEBiP/OsCERK1 receptor complex in at least 98.5% of 5232 surveyed rice accessions. These results demonstrate that *U. virens* utilizes a crucial virulence factor to subvert chitin-triggered flower immunity in most rice varieties, providing new insights into the susceptibility of rice to false smut disease.

**One Sentence Summary:** The fungal pathogen *Ustilaginoidea virens* disarms chitin-triggered immunity in rice flower via a secreted chitinase.

## INTRODUCTION

Flower-infecting fungal pathogens cause many detrimental crop diseases, such as Fusarium head blight in wheat (caused by *F. graminearum*) (Xu and Nicholson, 2009), Ergot disease in rye (caused by *Claviceps purpurea*) (Tudzynski and Scheffer, 2004), and corn smut disease (caused by *Ustilago maydis*) (Brefort et al., 2009). Some floral pathogens even introduce food toxins, such as DON produced by *F. graminearum* and Ergot alkaloids generated by *C. purpurea* (Tudzynski and Scheffer, 2004; Xu and Nicholson, 2009). *Ustilaginoidea virens* (Cooke) Takahashi (teleomorph: *Villosiclava virens*) is an emerging fungal pathogen infecting rice flower and causes rice false smut (RFS) disease, not only resulting in yield loss and quality reduction but also threatening the health of humans and animals due to *U. virens*-produced mycotoxins (Zhou et al., 2012; Sun et al., 2020). Numerous rice germplasms have been evaluated with different sensitivities to *U. virens* and a set of quantitative trait loci (QTL) for false smut field resistance have been mapped. However, neither fully-resistant rice cultivars have been identified nor false smut resistance genes have been cloned (Sun et al., 2020). Gene-for-gene resistance has not been found in rice against *U. virens*. Management and control of RFS disease will benefit from dissection of the compatible mechanism between rice flower and *U. virens*.

*U. virens* possesses specific infection strategies in rice flower. At late booting stage of rice, *U. virens* spores contacting rice developing spikelets can germinate on the surface of lemma and palea, on which no infection sites have been observed (Ashizawa et al., 2012; Tang et al., 2013). Instead, *U. virens* hyphae epiphytically extend into inner space of rice spikelets through the gap between the lemma and palea, and then primarily attack stamen filaments intercellularly (Ashizawa et al., 2012; Tang et al., 2013). *U. virens* can also infect lodicules, stigmas, and styles, but to a lesser extent (Tang et al., 2013; Song et al., 2016). After successful colonization, *U. virens* forms massive mycelia to embrace all the inner floral organs, and ultimately produces ball-shape fungal colonies named false smut balls, which are the only visible symptom of RFS disease (Fan et al., 2016; Sun et al., 2020). Although *U. virens* infects multiple floral parts, it requires rice stamens, but not pistils, for the formation of false smut balls (Fan et al., 2020). Again, *U. virens* infects roots and coleoptiles of rice, but cannot produce false smut balls in these organs (Ikegami, 1963; Schroud and TeBeest, 2005; Prakobsub and Ashizawa, 2017; Yong et al., 2018). It has been suggested that *U. virens* may hijack grain filling system in rice spikelets to obtain abundant nutrients for the formation of false smut balls (Fan et al., 2015; Song et al., 2016).

As a successful biotrophic pathogen (Zhang et al., 2014), first of all, *U. virens* should be able to evade or suppress host immunity. Previous transcriptome analyses indicate that expression of rice defense-related genes, such as *PAL* and *PR* genes, could be down-regulated upon *U. virens* infection (Fan et al., 2015; Han et al., 2015). *U. virens* can deploy a set of immunosuppressive effectors during infection (Zhang et al., 2014; Sun et al., 2020). Particularly, effector proteins such as SCRE1, UV_1261/SCRE2, and UV_5215 could inhibit cell death and/or pattern-triggered immunity (PTI) in plants. SCRE1 and UV_1261/SCRE2 both contribute to the virulence of *U. virens* in rice flower (Zhang et al., 2014; Fan et al., 2019; Fang et al., 2019; Zhang et al., 2020). However, their host targets and virulence mechanisms are unknown.

To counteract the infection of fungal pathogens, plants can mount a critical defense pathway called chitin-triggered immunity, i.e. chitin from the fungal cell wall is recognized by plant cell surface receptors to induce PTI (Jones and Dangl, 2006; Gong et al., 2020). In rice leaf organ, OsCEBiP functions as a major receptor with high affinity for chitin (Kaku et al., 2006; Hayafune et al., 2014). Two additional Lysin motif-containing proteins, OsLYP4 and OsLYP6, act as minor chitin receptors (Liu et al., 2012). As these chitin receptors lack a kinase domain, chitin signaling requires a co-receptor OsCERK1 to activate downstream signaling components (Shimizu et al., 2010). In turn, downstream cytoplasmic kinases such as OsRLCK118/176/185 regulate chitin-induced Ca^2+^ influx, activation of mitogen-activated protein kinase (MAPK), and burst of reactive oxygen species (ROS) (Wang et al., 2017; Fan et al., 2018; Wang et al., 2019). Compared to the well-documented leaf immunity in rice, little is known on the flower immunity. It is unrevealed whether OsCEBiP/OsCERK1-mediated chitin signaling is involved in the molecular interaction between rice flower and *U. virens*.

To understand the molecular events in the front line of battlefield between *U. virens* and rice flower, we previously performed a dual-transcriptome study on *U. virens* infecting with rice flower (Fan et al., 2015). Our subsequent studies focused on a number of *U. virens* genes whose transcriptional levels were highly increased during infection and the encoded proteins were putatively secreted (Fan et al., 2019). In this study, we report a candidate gene *UvChi1* that functions as a crucial virulence factor and subverts a highly conserved chitin-triggered immunity in rice flower. This study provides new insights into the pathogenic mechanism of *U. virens* and defense mechanism of rice flower, and gives implications for controlling rice false smut disease.

## RESULTS

### UvChi1 is a secreted protein essential for *U. virens* pathogenicity

In a previous transcriptome analysis, we found that *Uv5918* was among the most up-regulated genes in *U. virens* infecting rice flower (Fan et al., 2015). In this study, we determined a time-course expression pattern of *Uv5918* by RT-qPCR analysis. Compared to the expression level of *Uv5918* in axenic culture, the abundance of *Uv5918* transcripts started to increase at 5 day post inoculation (dpi), peaked at 7 dpi with more than 20-fold increase when *U. virens* hyphae invaded into the inner floral organs of rice spikelets (Fan et al., 2015) (Fig. 1A). Sequence analysis revealed that *Uv5918* (hereafter *UvChi1*) encoded a putative protein with 450 amino acid (aa), containing a predicted signal peptide (SP) and a putative chitinase active site (Supplemental Fig. S1). The secretion of UvChi1 was examined by Western blot analysis with an UvChi1-specific antibody. The control experiment with the GAPDH-specific antibody generated expected band only in the mycelia sample but not in the supernatant, indicating no contamination of fungal mass in the supernatant. By contrast, UvChi1 protein could be detected in both mycelia and supernatant fractions (Fig. 1B), indicating that UvChi1 could be secreted by *U. virens*. The functionality of UvChi1 SP was further verified by a yeast secretion assay as described previously (Jacobs et al., 1997; Fang et al., 2016; Fan et al., 2019). The sequence encoding predicted SP of UvChi1 was fused in frame with mature invertase (SUC2) and introduced into the yeast strain YTK12. The wild-type YTK12 cannot utilize raffinose due to its deficiency in invertase secretion, whereas the YTK12 strain transformed with the *UvChi1^SP^-SUC2* could grow well on YPRAA medium supplemented with raffinose as the sole carbon source. YTK12 strains transformed with *Avr1b^SP^-SUC2* or *Mg87 ^N-terminus^-SUC2* were used as the positive and negative control, respectively (Fig. 1C). As a result, UvChi1^SP^ is a functional SP.

**Figure 1.**
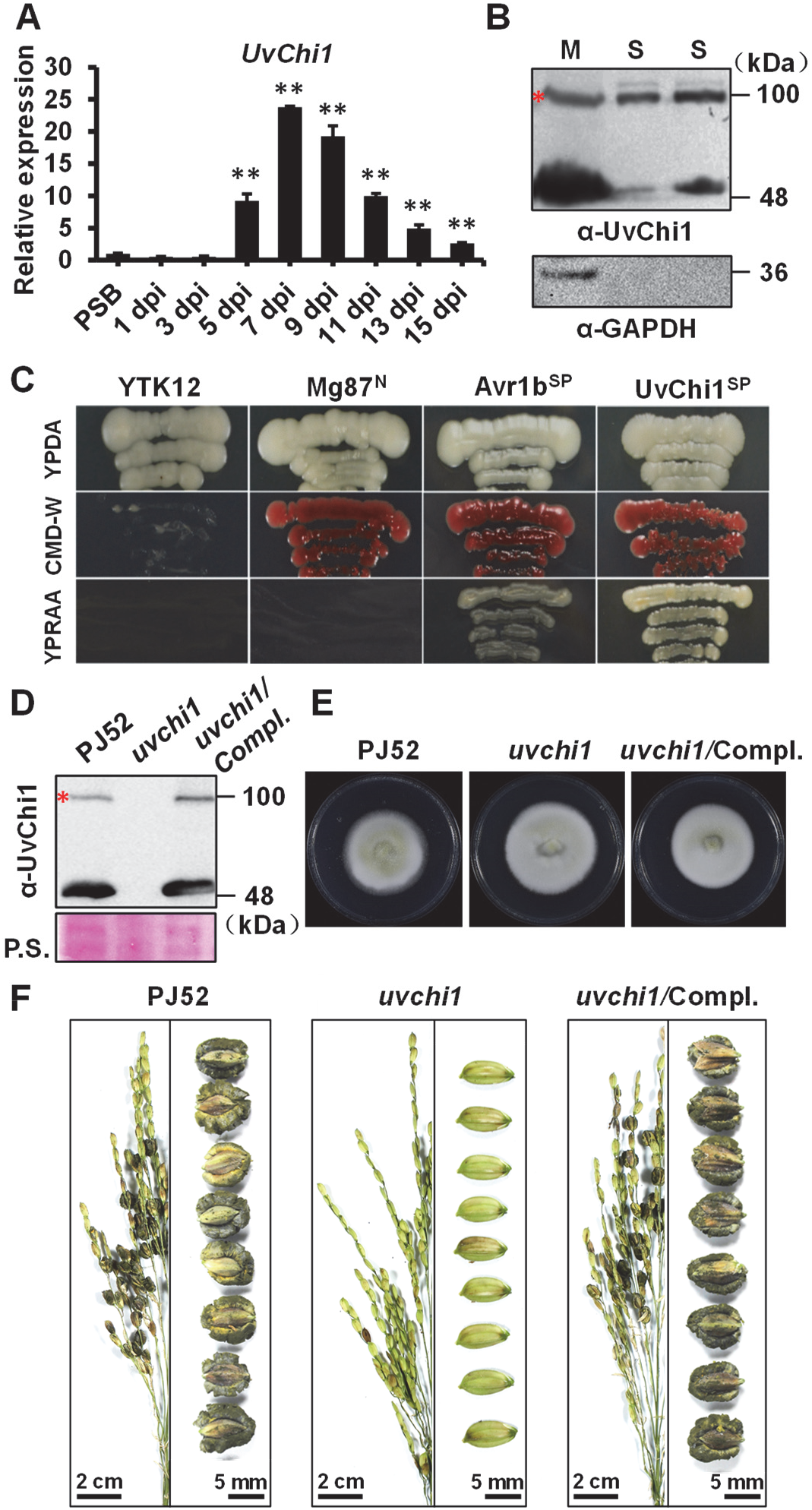
UvChi1 is a secreted protein essential for *Ustilaginoidea virens* pathogenicity in rice flower. **A,** Expression analysis of *UvChi1* during *U. virens* infection of rice. Spikelets from PJ52-inoculated rice panicles were sampled at indicated time points and subjected to RT-qPCR analysis. The mixture of mycelia and conidia from PSB-cultured PJ52 was collected as the control sample. Relative expression level of *UvChi1* was determined using *UvTub2α* as the reference gene. Data are represented as means ± SD of three biological replicates. Asterisk indicates significant difference determined by Student’s *t* test (\**P* < 0.05, ** *P* < 0.01). Similar results were obtained from two independent experiments. dpi, day post inoculation. **B,** UvChi1 can be secreted into the culture medium. Total protein from PSB-cultured *U. virens* mycelia and cultured supernatants were subjected to Western blot analysis. Red asterisk indicates putative dimer of UvChi1. α-UvChi1, anti-UvChi1 antibody. α-GAPDH, anti-glyceraldehyde-3-phosphate dehydrogenase antibody. M, mycelia. S, supernatant. **C,** Validation of UvChi1 signal peptide (SP). The DNA fragment encoding SP of UvChi1 was cloned into pSUC2, in frame with an invertase gene. The resultant plasmid was transformed into YTK12 that is unable to utilize raffinose. SP of UvChi1 could enable YTK12 to grow on YPRAA medium, indicating its functionality. SP of Avr1b and N terminus of Mg87 were applied as positive and negative controls, respectively. **D,** Western blot analysis of *UvChi1* knockout and complementation strains using UvChi1-specific antibody. **E,** Top view of *UvChi1* knockout and complementation strains, and wild-type PJ52 cultured in PSA media for two weeks. **F,** Pathogenicity assay of *UvChi1* knockout and complementation strains, and PJ52. Inocula of indicated *U. virens* strains were injected into the panicles (*n* ≥ 30 for each strain) of rice accession Q455 at late booting stage. Disease phenotype was recorded at four week post inoculation (wpi). Note that no false smut balls were formed in *uvchi1* mutant-inoculated rice panicles.

To determine the role of *UvChi1* in pathogenicity, we knockout it in *U. virens* using a CRISPR-Cas9-assisted gene replacement approach (Liang et al., 2018), and obtained multiple knockout mutants (Supplemental Fig. S2). Markedly, knockout mutant *uvchi1* lost pathogenicity in rice panicles, i.e. failing to develop RFS balls. The complementation strain could restore the ability to form RFS balls (Fig. 1D, F; Supplemental Fig. S2). By contrast, the *uvchi1* knockout mutant showed normal colony morphology comparable to wild-type and complementation strains (Fig. 1E). These data suggest that *UvChi1* is an essential virulence factor of *U. virens*.

### UvChi1 suppresses chitin-induced immunity in rice

To explore the virulence mechanism of UvChi1, we first assessed whether UvChi1 modulate immune response in rice. We generated transgenic rice ectopically over-expressing *UvChi1*and confirmed its expression in both leaf and flower organs (Fig. 2). In both leaf and flower organs of wild-type rice, chitin could induce the expression of defense-related genes, such as *OsBETV1*, *OsNAC4*, and *OsPR10b*. Markedly, the induction of these genes was suppressed in UvChi1-expressing leaves and flowers (Fig. 2), supporting a role of UvChi1 in blocking rice immunity to promote infection (Supplemental Fig. S3).

**Figure 2.**
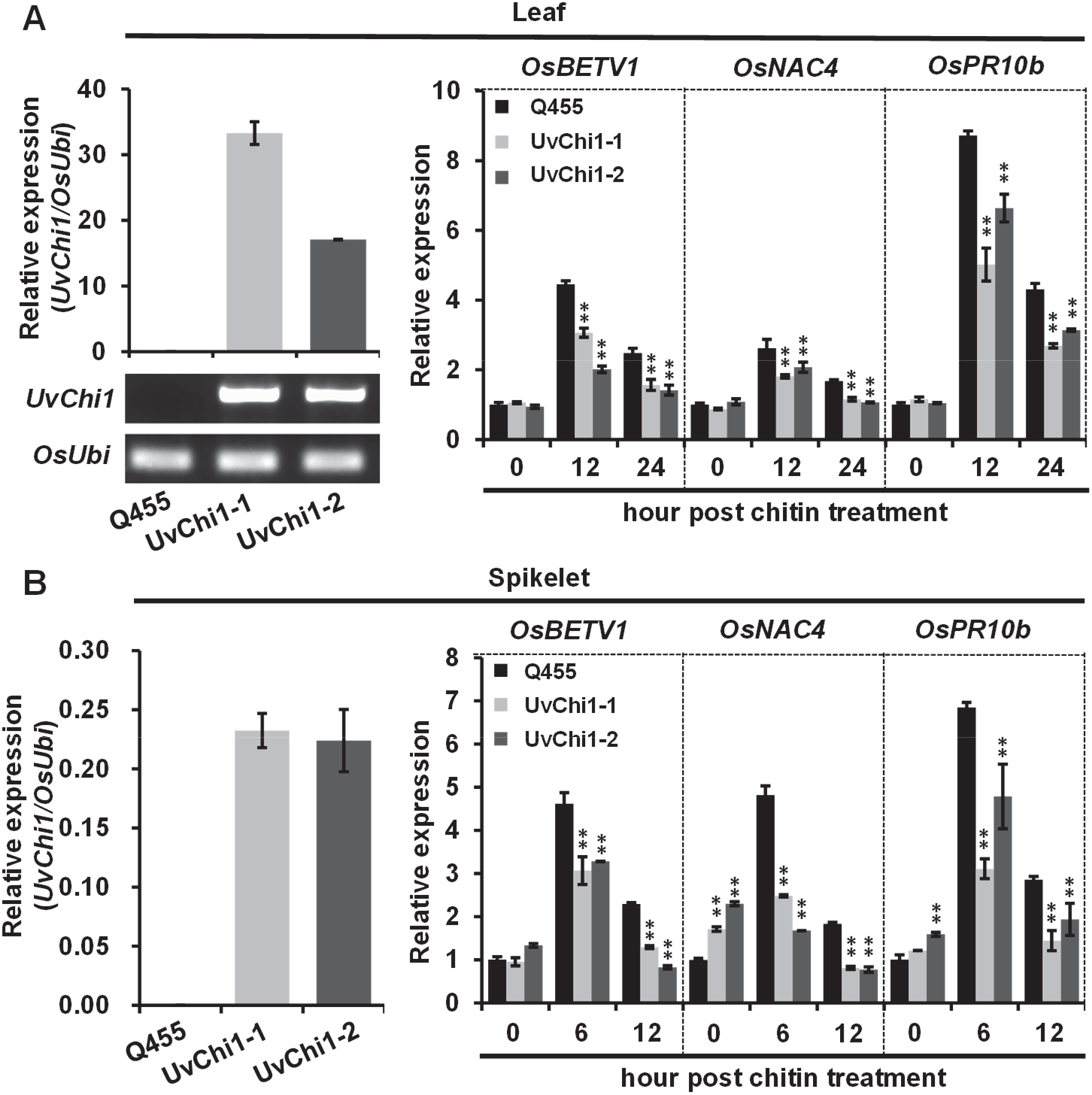
Ectopic expression of *UvChi1* suppresses chitin-induced expression of defense-related genes in rice. *UvChi1* was driven by a 35S promoter and expressed in rice accession Q455. The expression of *UvChi1* was confirmed in leaf (**A**) and spikelet (**B**) of transgenic lines by RT-qPCR. Then, leaf and spikelet samples were respectively treated with chitin and sampled at indicated time points for RT-qPCR analysis of rice defense-related genes. Relative expression levels of indicated genes were determined using *OsUbi* as the reference gene. Data are represented as means ± SD of three biological replicates. Asterisk indicates significant difference determined by Student’s *t* test (** *P* < 0.01).

As reported, fungal chitinases have evolved as effector proteins to prevent chitin-triggered plant immunity via their ability of binding with chitin or degrading chitin oligomers (Fiorin et al., 2018; Han et al., 2019; Yang et al., 2019; Martínez-Cruz et al., 2021). In consistent with these reports, *UvChi1* encoded an enzymatically active fungal chitinase (Supplemental Fig. S1), which possessed chitin-binding ability and could suppress chitin-triggered ROS burst and induction of defense gene expression in rice (Supplemental Fig. S4).

### UvChi1 interacts with OsCEBiP and OsCERK1 to impair their mediated chitin signaling in rice

Fungal chitinase effectors may target to or be recognized by plant cell surface proteins, which is supported by that *Magnaporthe oryzae* MoChia1 can interact with rice plasma membrane proteins OsMBL1 and OsTPR1 (Han et al., 2019; Yang et al., 2019). OsMBL1 is a jacalin-related Mannose-Binding Lectin protein contributing to chitin perception and chitin-triggered rice immunity, which can be suppressed by MoChia1 (Han et al., 2019). OsTPR1 is a tetratricopeptide-repeat family protein and functions as an immune receptor recognizing MoChia1 to regain chitin-triggered immunity in rice (Yang et al., 2019). We found that, unlike MoChia1, UvChi1 could not interact with OsMBL1 and OsTPR1 (Supplemental Fig. S5), although *OsMBL1* and *OsTPR1* were highly expressed in rice flower organ (Supplemental Fig. S6).

Cell surface receptors OsCEBiP and OsCERK1 play central roles in chitin-triggered immunity in rice (Gong et al., 2020). As *OsCEBiP* and *OsCERK1* were expressed even higher in rice flowers than in leaves (Supplemental Fig. S6), we tested whether OsCEBiP and OsCERK1 could interact with UvChi1. In GST pull-down assay, OsCEBiP or OsCERK1 was expressed with a GST tag, and UvChi1 was expressed with an MBP tag. MBP-UvChi1 could be pull-down by both GST-OsCEBiP and GST-OsCERK1, but not by GST alone, indicating direct interactions of UvChi1 with OsCEBiP and OsCERK1 (Fig. 3A). Co-IP assay further confirmed that UvChi1 was physically associated with OsCEBiP and OsCERK1 (Fig. 3B). Interestingly, UvChi1 protein mutated at its putative chitin-binding sites (UvChi1^mcb^) lost chitin-binding/degrading ability (Supplemental Fig. S7), but could still interact with OsCEBiP and OsCERK1 both *in vitro* and *in vivo* (Fig. 3).

**Figure 3.**
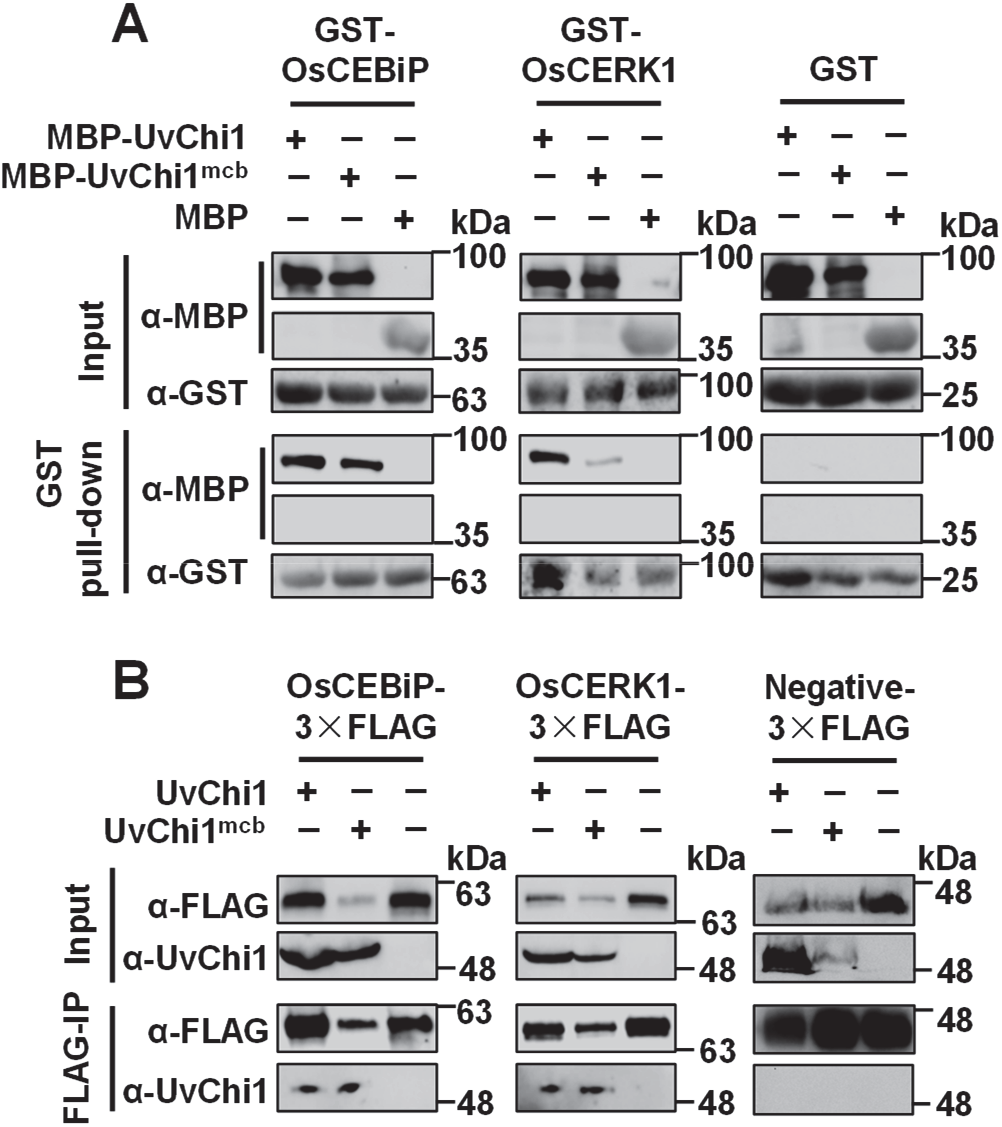
UvChi1 interacts with OsCEBiP and OsCERK1. **A,** *In vitro* GST pull-down assay. The recombinant proteins GST-OsCEBiP, GST-OsCERK1, MBP-UvChi1, and MBP-UvChi1^mcb^ (mutated at chitin binding sites) were purified from *Escherichia coli*. GST and MBP tag proteins were used as negative controls. Protein interaction was visualized with Western blot. **B,** *In vivo* co-immunoprecipitation (Co-IP) assay. OsCEBiP-3×FLAG or OsCERK1-3×FLAG was co-expressed with UvChi1, UvChi1^mcb^ in *Nicotiana benthamiana*. An FLAG-tagged protein without interaction with UvChi1 was served as the negative control. IP was conducted using anti-FLAG affinity gel and subjected to Western blot analysis using anti-FLAG or anti-UvChi1 antibodies. Note that the coding region of *OsCEBiP* and *OsCERK1* were amplified from NPB, representing polymorphic type 1 (T1) as depicted in Figure 5.

Chitin-induced oligomerization of chitin receptors is a prerequisite for intracellular chitin signaling in Arabidopsis and rice (Gong et al., 2020). We thus tested whether UvChi1 could affect the oligomerization of OsCERK1 and OsCEBiP. We conducted Co-IP assays in *Nicotiana benthamiana*, and found that UvChi1 could reduce the interactions of OsCEBiP-OsCEBiP, OsCEBiP-OsCERK1, and OsCERK1-OsCERK1 (Fig. 4A). The competition effects became stronger along with the increasing amounts of UvChi1 protein (Fig. 4A). Similar results were obtained for UvChi1^mcb^. These results suggest that UvChi1 disrupts homo- and hetero-oligomerization of OsCERK1 and OsCEBiP, which is independent on its chitin-binding sites.

**Figure 4.**
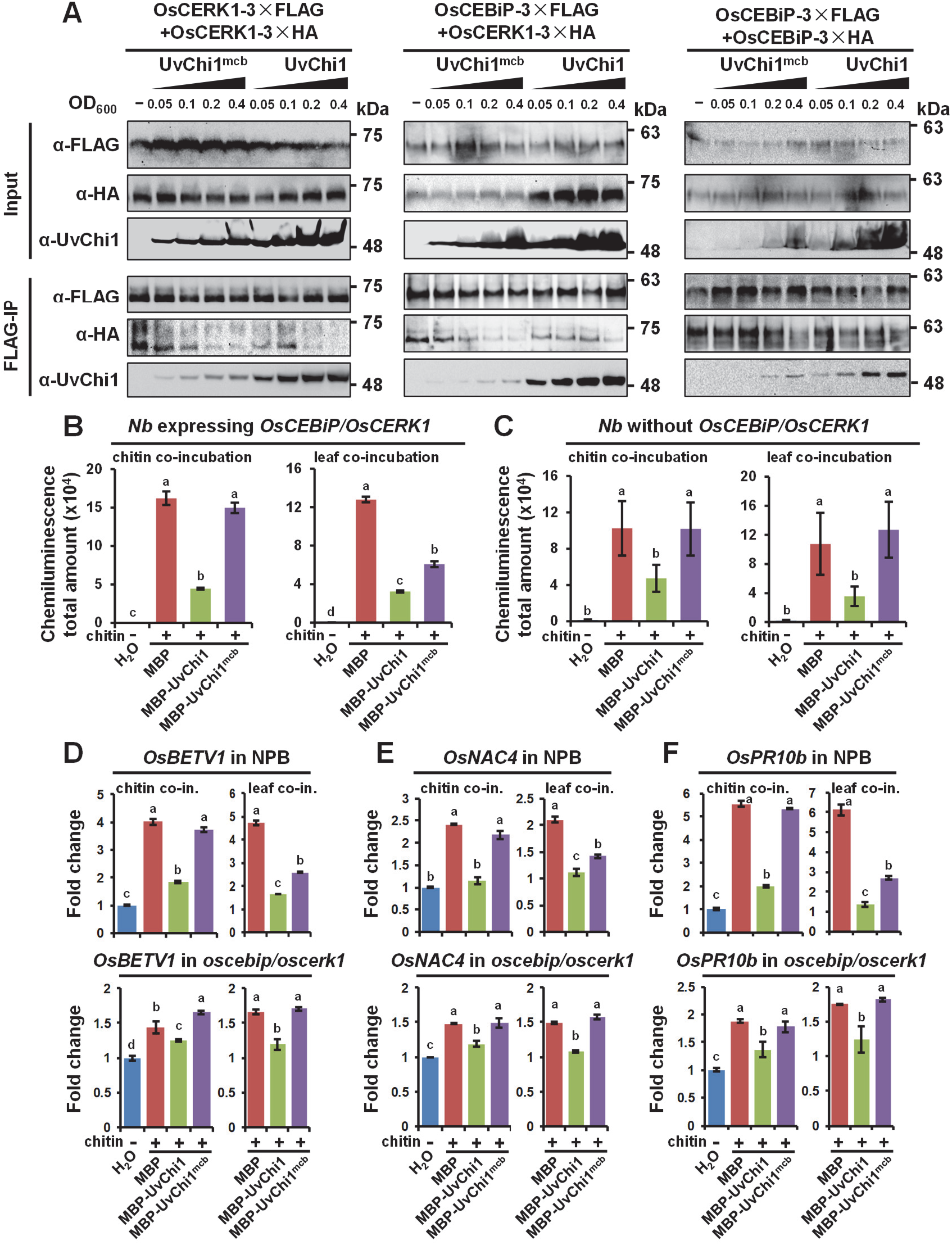
UvChi1 impairs OsCEBiP/OsCERK1-mediated chitin signaling in rice. **A,** UvChi1 reduces homo- and hetero-oligomerizations of OsCERK1 and OsCEBiP. For competitive Co-IP assay, OsCERK1 and OsCEBiP tagged with FLAG or HA was transiently co-expressed with/without UvChi1 or UvChi1^mcb^ in *Nicotiana benthamiana*. At 48 hour post infiltration, leaves were treated with chitin by infiltration for 10 min prior to protein extraction. IP was conducted using anti-FLAG affinity gel and subjected to Western blot analysis using anti-FLAG, anti-HA, or anti-UvChi1 antibodies. **B,** ROS assay in *Nicotiana benthamiana* (*Nb*) transiently expressing *OsCEBiP* and *OsCERK1*. *Nb* leaves were infiltrated with GV3101 strains harboring expression constructs of *OsCEBiP* and *OsCERK1*. Leaf discs were sampled for ROS assay at 36 hour post infiltration. The indicated recombinant proteins were either co-incubated with chitin for 1 h (namely chitin co-incubation treatment) before ROS detection, or co-incubated with leaf discs for 6 h (namely leaf co-incubation treatment) before measurements of chitin-induced ROS. Please refer to Materials and Methods for details. Data are represented as mean ± SD of four biological replicates. Different letters above data bars indicate significant difference as determined by one-way ANOVA with post hoc Tukey HSD analysis (*P* < 0.05). Similar results were obtained from two independent experiments. **C,** ROS assay in *Nb* leaves without expressing *OsCEBiP* and *OsCERK1*. Measurements of ROS and data analysis were the same as in **B**. Note that the coding region of *OsCEBiP* and *OsCERK1* were amplified from NPB, representing polymorphic type 1 (T1) as depicted in Figure 5. **D-F,** Expression analysis of defense-related genes in rice leaves of NPB and *oscebip*/*oscerk1* double mutant. Leaf discs were sampled at 4-leaf stage rice seedlings. The indicated recombinant proteins were either co-incubated with chitin or co-incubated with leaf discs before analysis of chitin-induced gene expression. Data are represented as mean ± SD of three repeats. Different letters above data bars indicate significant difference as determined by one-way ANOVA with post hoc Tukey HSD analysis. Similar results were obtained from three independent biological experiments.

**Figure 5.**
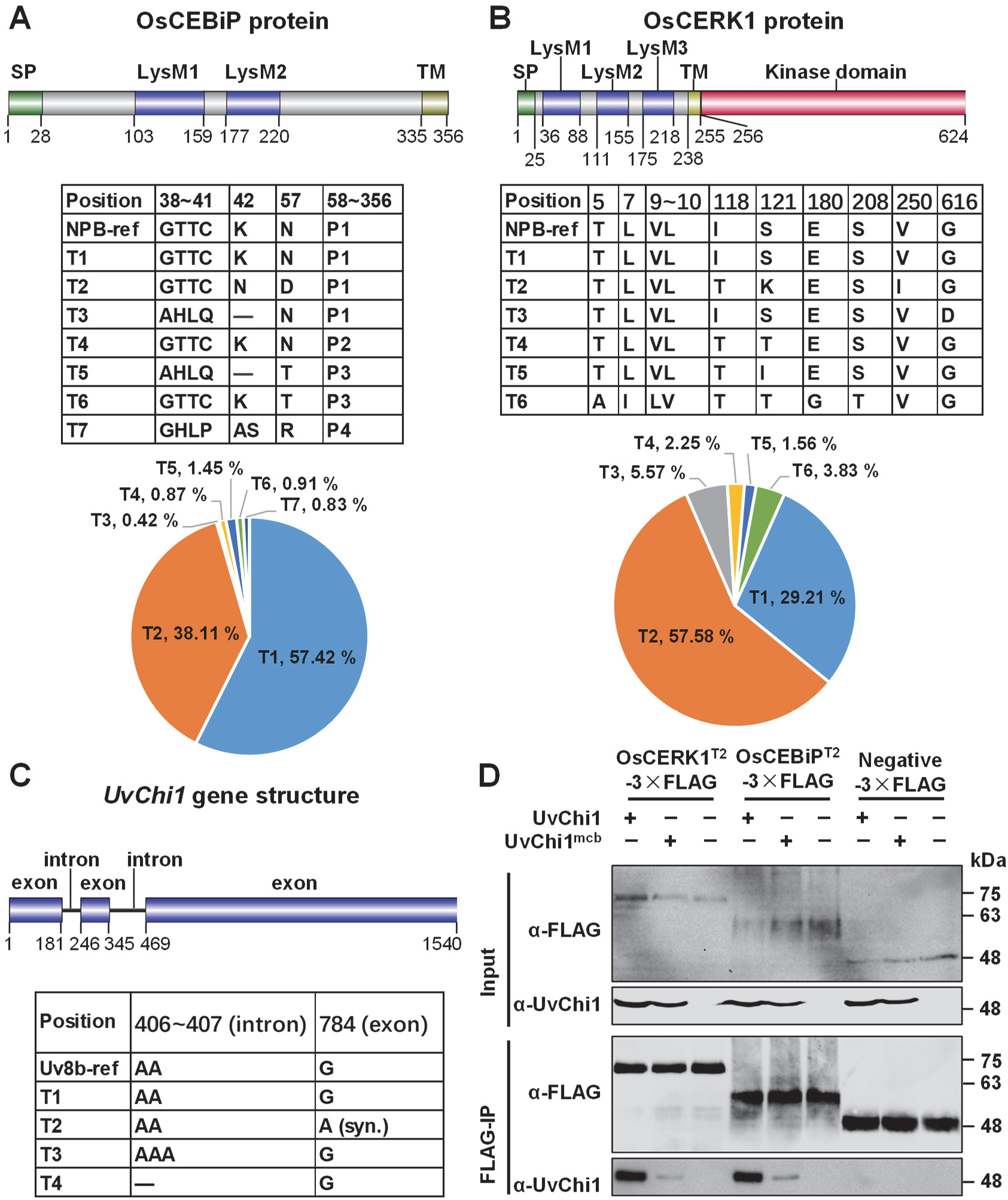
UvChi1-OsCEBiP/OsCERK1 interaction module is highly conserved in the *Ustilaginoidea virens*-rice pathosystem. **A-B,** Polymorphism analysis of OsCEBiP **(A)** and OsCERK1 **(B)** protein sequences. Schematic diagram of protein domains was shown. Protein sequences of OsCEBiP and OsCERK1 in over 5000 rice accessions were retrieved from MBKbase-rice database (http://mbkbase.org/rice). Polymorphic types (T1-T7) were determined using Nipponbare as the reference. Percentage for each polymorphic type was shown as pie charts. Protein sequences representing P1-P4 were presented in Supplemental Fig. S9. SP, signal peptide. LysM, lysin motif. TM, transmembrane domain. **C,** Polymorphism analysis of UvChi1. Schematic diagram of *UvChi1* gene structure was displayed. Genomic sequences of *UvChi1* from over 50 *U. virens* isolates originated from North, West, Middle, and East China, as well as Japan and Nepal, were amplified and sequenced. Polymorphic types (T1-T4) were determined using Uv8b strain as the reference. One InDel in an intron and one synonymous substitution (syn.) in an exon were detected. **D,** Co-IP assay. As polymorphic type T1 (Nipponbare reference type) plus T2 of OsCEBiP and OsCERK1 accounted for 86.79%-95.53% in over 5000 rice accessions. OsCEBiP^T2^ and OsCERK1^T2^ were also tested for interactions with UvChi1. OsCEBiP^T2^-3×FLAG or OsCERK1 ^T2^-3×FLAG was co-expressed with UvChi1 or UvChi1^mcb^ in *Nicotiana benthamiana*. An FLAG-tagged protein without interaction with UvChi1 was served as the negative control. IP was conducted using anti-FLAG affinity gel and subjected to Western blot analysis using anti-FLAG or anti-UvChi1 antibodies.

Next, we intended to assess whether OsCEBiP/OsCERK1-mediated immune responses were suppressed by UvChi1. To exclude the immunosuppressive effects of UvChi1 resulting from its chitin-binding/degrading ability, we included UvChi1^mcb^ in the subsequent experiments. In *N. benthamiana* expressing *OsCEBiP*/*OsCERK1*, chitin-induced ROS production could be suppressed by MBP-UvChi1 but not by MBP-UvChi1^mcb^ and MBP, when chitin was pre-incubated with the recombinant proteins. However, when leaves were pre-incubated with the recombinant proteins before chitin treatment (which may give enough time for the recombinant proteins to approach the cell surface receptors OsCEBiP and OsCERK1), chitin-induced ROS could be inhibited by both MBP-UvChi1 and MBP-UvChi1^mcb^ (Fig. 4B). In wild-type rice NPB, chitin-induced expression of defense-related genes could be repressed by MBP-UvChi1 but not by MBP-UvChi1^mcb^ when chitin was pre-incubated with the recombinant proteins; whilst MBP-UvChi1^mcb^ showed markedly immunosuppressive effects when it was pre-incubated with leaf discs before chitin treatment (Fig. 4D-F). By contrast, in *N. benthamiana* without expression of *OsCEBiP*/*OsCERK1* and in rice *oscebip*/*oscerk1* double mutant (Supplemental Fig. S8), pre-incubation of leaves with MBP-UvChi1^mcb^ no longer blocked chitin-induced ROS and expression of defense-related genes (Fig. 4C-F). These data indicate that UvChi1^mcb^ can suppress chitin-triggered plant immunity in an OsCEBiP/OsCERK1-dependent manner.

### UvChi1 targeting OsCEBiP/OsCERK1 is highly conserved in the *U. virens*-rice pathosystem

We intended to test whether the interaction module of UvChi1–OsCEBiP/OsCERK1 was conserved in the *U. virens*-rice pathosystem. We first analyzed the polymorphisms of OsCEBiP and OsCERK1 protein sequences among over 5000 rice accessions, of which the genomic data are retrieved from the database of MBKbase-rice (http://mbkbase.org/rice) (Peng et al., 2020). We detected seven and six polymorphism types for OsCEBiP and OsCERK1, respectively. For OsCEBiP, the majority (57.42%) of 4962 detected rice accessions had the identical protein sequence to the reference NPB; 38.11% (designated as polymorphism type 2, T2) had two amino acid changes located at position 42 and 57; 4.06% had truncations at the C-terminus of OsCEBiP (Fig. 5A; Supplemental Fig. S9). For OsCERK1, 29.21% of 5066 detected rice accessions possessed identical protein sequence to NPB; 57.58% (designated as T2) had three amino acid differences at position 118, 121, and 250 (Fig. 5B). We then analyzed the polymorphism of UvChi1 among over 50 *U. virens* field isolates originated from different rice production areas in China, as well as in Japan and Nepal (Supplemental Fig. S11). Surprisingly, we detected no polymorphisms for UvChi1 protein sequences from all the tested *U. virens* isolates, although one InDel was detected in an intron and one synonymous substitution was found in an exon of *UvChi1* gene (Fig. 5C).

Since OsCEBiP^NPB^ and OsCERK1^NPB^ interacted with UvChi1 (Fig. 3), we tested the interactions between OsCEBiP^T2^ (OsCERK1^T2^) and UvChi1. Interestingly, both OsCEBiP^T2^ and OsCERK1^T2^ could interact with UvChi1 and UvChi1^mcb^ (Fig. 5D), and oligomerizations of OsCEBiP^T2^ and OsCERK1^T2^ were also disrupted by UvChi1 and UvChi1^mcb^ (Supplemental Fig. S10). These data indicate that UvChi1 can target OsCEBiP in at least 95.53% of 4962 rice accessions, and OsCERK1 in at least 86.79% of 5066 accessions. Collectively, highly conserved UvChi1 interferes with either OsCEBiP or OsCERK1 in over 98.5% of 5232 rice accessions.

### UvChi1 disarms OsCEBiP/OsCERK1-mediated resistance to *U. virens*

To assess the role of OsCEBiP/OsCERK1 in rice resistance to *U. virens*, we inoculated the *U. virens* PJ52 and *uvchi1* mutant into wild-type rice and *oscebip*/*oscerk1* double mutant (Supplemental Fig. S8). When compared to the high virulence of PJ52, the *uvchi1* mutant lost ability of developing false smut balls in both WT and *oscebip*/*oscerk1* plants (Fig. 6A, E). We further checked the infection process of *uvchi1* mutant in rice panicles, and surprisingly found that some spikelets were indeed infected by *uvchi1*, of which the fungal mycelia embraced the inner floral organs including stamens and pistils (Fig. 6D). This infection status has been reported to be a prerequisite for symptom development of false smut disease (Fan et al., 2020). We quantified the numbers of infected panicle and infected spikelet per panicle, and observed that the reduced virulence caused by deletion of *UvChi1* was relieved in *oscebip*/*oscerk1* mutant compared with WT (Fig. 6B, C, F, G). These data suggest that OsCEBiP/OsCERK1 contributes to rice resistance to the deletion mutant *uvchi1*; nevertheless, UvChi1 can disarm this immune pathway.

**Figure 6.**
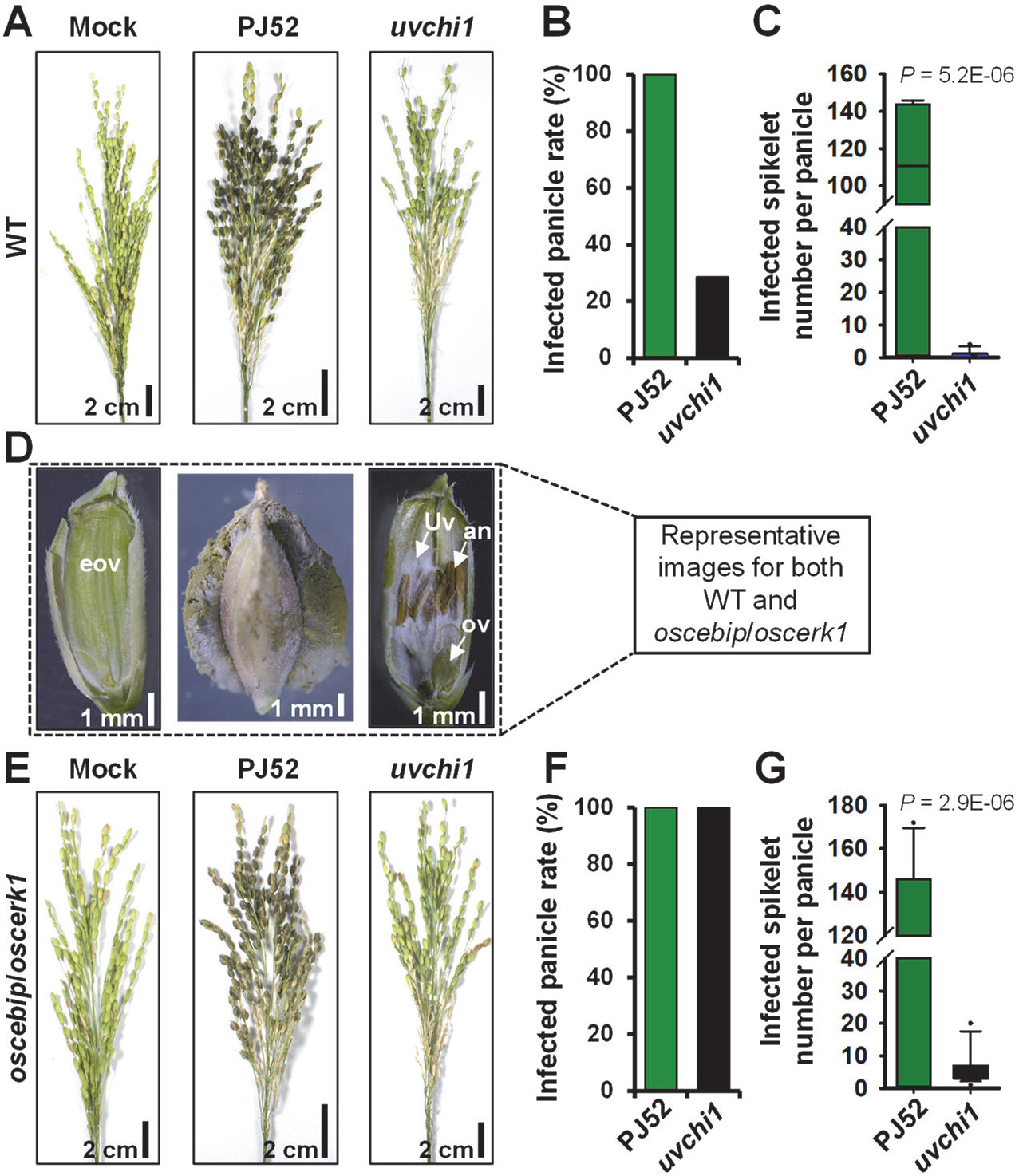
The role of OsCEBiP/OsCERK1 in resistance to *Ustilaginoidea virens*. Disease assay of wide-type (WT) Q455 **(A-C)** and *oscebip*/*oscerk1* double mutant **(E-G)** infected with *U. virens* PJ52 and *uvchi1* knockout mutant. Note that *uvchi1* mutant failed to form false smut balls in either WT or *oscebip*/*oscerk1*, but could infect rice spikelets to some extent, i.e. *U. virens* mycelia embracing the inner floral organs **(D)**. The infected panicle rate **(B, F)** and infected spikelet number per diseased panicle **(C, G)** were quantified. Data of infected spikelet number are box-plotted (*n* > 15). *P* values were determined by Student’s *t* test. an, anther. Uv, *U. virens*. ov, ovary. eov, expanded ovary.

## DISCUSSION

Chitin-triggered immunity is a conserved anti-fungal immune system in plants. CEBiP/CERK1-mediated chitin signaling is a central plant immunity against leaf-infecting fungal pathogens (Liu and Wang, 2016; Gong et al., 2020), whilst this chitin signaling pathway was yet to be verified in plant flower organ. In the present work, we first showed that *OsCEBiP* and *OsCERK1* were highly expressed in rice spikelets (Supplemental Fig. S6); second, chitin treatment of rice spikelets could induce the expression of PTI marker genes, such as *OsNAC4*, *OsPR10b*, and *OsBETV1*, which was similarly observed in rice leaves (Fig. 2; Supplemental Fig. S4); third, chitin-induced expression of PTI marker genes was markedly attenuated in the spikelets expressing or treated with UvChi1 (Fig. 2; Supplemental Fig. S4); fourth, *oscebip*/*oscerk1* mutant had enhanced susceptibility to the *U. virens uvchi1* (Fig. 6). These data support a functional OsCEBiP/OsCERK1-mediated immunity in rice flower organ.

Chitin-triggered immunity can be subverted by sophisticated fungal pathogens via multiple strategies as summarized in a recent review (Gong et al., 2020). For instance, fungal pathogens protect their cell wall from releasing plant immunity-inducible chitin fragments by secreting proteases to degrade plant chitinases (Okmen et al., 2018), masking cell wall with α-1,3-glucan and effector proteins such as Avr4 and Avr4-likes (van den Burg et al., 2006; Marshall et al., 2011; Fujikawa et al., 2012), or deacetylating chitin to chitosan (Gao et al., 2019). Fungal pathogens can also secrete effectors with high affinity to chitin to outcompete host chitin receptors, and/or with ability of degrading chitin oligomers, such as Ecp6 (de Jonge et al., 2010), Slp1 (Mentlak et al., 2012), MoAa91 (Li et al., 2020), MoChia1 (Han et al., 2019; Yang et al., 2019), and EWCAs (Martínez-Cruz et al., 2021). Some fungi can utilize effectors like NIS1 and AvrPiz-t to target plant intracellular immune components involved in chitin signaling, such as BIK1 and OsRac1 (Bai et al., 2019; Irieda et al., 2019). In this study, we found that a secreted chitinase UvChi1 from *U. virens* directly targeted to the chitin receptor OsCEBiP and the co-receptor OsCERK1 to interfere with their mediated chitin signaling, in addition to the MoChia1-like roles of outcompeting rice receptors for chitin binding and degrading chitin (Supplemental Figs. S1, S4, S11) (Han et al., 2019). Our findings add new knowledge to the fungal counterstrategies for dampening chitin-triggered plant immunity, especially in floral organ.

Fungal chitinases have versatile roles in fungal nutrition, autolysis, growth, and development (Hartl et al., 2012). Recent studies uncover novel functions of fungal chitinases in manipulating plant immunity. *Moniliophthora perniciosa*, *M. roreri M. oryzae*, and *Podosphaera xanthii* secrete enzymatically inactive or active chitinases as virulence factors to sequester chitin-triggered host immunity (Fiorin et al., 2018; Han et al., 2019; Yang et al., 2019; Martínez-Cruz et al., 2021). Accordingly, plant hosts have evolved cell-surface receptors to fight against these chitinase effectors. For example, rice utilizes the plasma membrane-localized OsTPR1 to recognize chitinase effector MoChia1 from a leaf-infecting pathogen *M. oryzae*, thereby releasing free chitin to re-trigger chitin-induced plant immunity (Yang et al., 2019). In our work, UvChi1 could not interact with OsTPR1 (Supplemental Fig. S5), thus possibly escaping from the surveillance of OsTPR1-activated immunity. Whether OsTPR1 can recognize other chitinase family members in *U. virens* needs to be clarified in future. Another plasma membrane-localized protein OsMBL1 can compete with the chitinase effector MoChia1 on chitin binding and cause earlier induction of defense response in rice cells, likely serving as a novel chitin sensor (Han et al., 2019). As OsMBL1 could not interact with UvChi1 (Supplemental Fig. S5) and *OsMBL1* gene was highly expressed in rice flowers (Supplemental Fig. S6), chitin-OsMBL1 signaling may function in rice flower immunity against *U. virens*. It could not be ruled out that the minor chitin receptors LYP4 and LYP6 (Liu et al., 2012) may be also involved in rice resistance to *U. virens*. Moreover, rice genome possesses a large pool of cell surface receptor-encoding genes, such as >1000 *RLKs* and 90 *RLPs* (Shiu et al., 2004; Fritz-Laylin et al., 2005). It would be interesting to identify candidate receptors that recognize *U. virens*-derived PAMPs/elicitors, and to characterize their roles in rice resistance to *U. virens* (Supplemental Fig. S11).

Notably, we observed that the deletion mutant *uvchi1* was unable to develop false smut balls in *oscebip*/*oscerk1* double mutant as well as in WT, although the infection rate increased to some extent in *oscebip*/*oscerk1* compared to that in WT (Fig. 6). This suggests that UvChi1 may function more than modulating chitin perception and signaling in rice. UvChi1 can possibly target other plant immune components to promote *U. virens* infection, or manipulate rice metabolisms to gain abundant nutrients for the formation of false smut balls. Identifying other host targets of UvChi1 will help to unveil novel virulence mechanisms of *U. virens*.

Sequence analysis revealed that OsCEBiP and OsCERK1 were highly conserved among over 5000 rice accessions, suggesting a conserved and central role of chitin-OsCEBiP/OsCERK1 signaling in anti-fungal immunity of rice. Nevertheless, this immune pathway was dampened by *U. virens* via UvChi1 (Fig. 6), of which the protein sequence had no variations among all the examined *U. virens* isolates (Fig. 5C). In conclusion, *U. virens* can deploy a core effector to subvert a conserved anti-fungal immunity in rice, which could well-explain why *U. virens* is generally compatible with rice. Importantly, as UvChi1 is essential for *U. virens* pathogenicity to develop false smut balls (Fig. 1F and Fig. 6), it may serve as a promising target for the development of novel effective fungicides against *U. virens*.

## MATERIALS AND METHODS

### Fungal and plant materials, growth conditions, and disease assay

A virulent *U. virens* strain PJ52-2-5 (PJ52 for short) (Wang et al., 2016) was used in this work. It was isolated from a false smut ball naturally formed in rice accession Pujiang 6 in Sichuan, China. PJ52 was stored as mycelial clumps at −80°C, and was reactivated on potato sucrose agar (PSA) at 28°C before use. Rice accession Nipponbare (NPB) and a germplasm Q455 were used in this study. For RT-qPCR and reactive oxygen species assays, rice plants were grown in climatic chambers at 28°C, 14 h light /25°C, 10 h darkness. For rice false smut disease assay, rice plants were grown in an experimental field under natural conditions, and were maintained without spraying any fungicides at all growth stages of rice.

Infection of rice with *U. virens* was performed as described previously with minor modifications (Fan et al., 2015). Briefly, the potato sucrose broth (PSB)-cultured mixture of mycelia and conidia of *U. virens* strains were collected as inoculum, which was artificially injected with a syringe into rice panicles at late booting stages. Disease symptoms were photographed and disease severity was recorded at around four week post inoculation (wpi).

### Constructs and transformation

For validation of UvChi1 signal peptide (SP), predicted *UvChi1^SP^* sequence was synthesized by Sangon Biotech (Chengdu, China) and cloned into the pSUC2T7M13ORI (pSUC2) vector (Jacobs et al., 1997) to generate pSUC2-UvChi1^SP^.

To generate gene knockout and complementation plasmids for *UvChi1*, 955-bp upstream and 1034-bp downstream sequences were amplified and subcloned into a gene replacement vector pRF-HU2 (Frandsen et al., 2012) to generate pRF-HU2-UvChi1. To improve homologous recombination efficiency in *U. virens* gene knockout experiments, pCas9-tRp-gRNA-UvChi1 plasmid was constructed by introducing a *UvChi1*-specific gRNA spacer into the pCas9-tRp-gRNA vector following previous reports (Liang et al., 2018; Guo et al., 2019). The hygromycin-resistant gene *Hph* in pSK1044 vector (Yu et al., 2015) was replaced by a basta-resistant gene *bar* amplified from the vector Pzp-Bar-Ex (Fan et al., 2019), resulting in vector SK1044-Bar. The entire *UvChi1* gene including 2.0-kb native promoter sequence and 0.5-kb downstream sequence was amplified and ligated into the *Eco*RI-*Xho*I linearized SK1044-Bar vector. The primers and gRNA spacer sequences are listed in Supplemental Table S2.

For purification of recombinant proteins, coding sequences of *UvChi1*, *UvChi1^mcb^*, *OsCEBiP*, *OsCERK1*, *OsMBL1*, *OsTPR1*, and *MoChia1* were amplified with indicated primers (Supplemental Table S2) and ligated into the *Bam*HI-*Eco*RI linearized pMAL-c5x or pGEX-6p-1 vectors. *UvChi1^mcb^* was obtained through mutating the chitin binding sites in *UvChi1*.

To generate constructs for Co-IP experiments, the coding sequences of *UvChi1* or *UvChi1^mcb^* were amplified and cloned into the pCAMBIA1300 vector to generate plasmids 35S-UvChi1 and 35S-UvChi1^mcb^. The coding sequences of *OsCEBiP* or *OsCERK1* were amplified and cloned into the pCAMBIA1300-3×FLAG or pCAMBIA1300-3×HA vectors to generate plasmids OsCEBiP-3×FLAG, OsCERK1-3×FLAG, OsCEBiP-3×HA, and OsCERK1-3×HA. OsCEBiP^T2^-3×FLAG, OsCERK1^T2^-3×FLAG, OsCEBiP^T2^-3×HA, and OsCERK1^T2^-3×HA plasmids were obtained by mutating polymorphic sites to type 2 (T2) as indicated in Fig. 5. The primer sequences and related information are presented in Supplemental Table S2.

To make CRISPR-Cas9 constructs for simultaneously knocking-out *OsCEBiP* and *OsCERK1*, the gene-specific guide RNAs were designed with an online software toolkit CRISPR-GE (Xie et al., 2017) and subcloned into the pRGEB32 binary vector, resulting in Cas9-OsCEBiP/OsCERK1 construct. Guide RNA sequences and primers are listed in Supplemental Table S2.

To knockout *UvChi1* in *U. virens*, gene replacement construct pRF-HU2-UvChi1 and pCas9-tRP-gRNA-UvChi1 were co-transformed into protoplasts of *U. virens* strain PJ52 as described by a previous study (Talbot et al., 1993). Knockout mutants were screened from hygromycin-resistant transformants by PCR. The positions of primers are indicated in Fig. S2. For complementation assay, the construct SK1044-Bar-UvChi1 (Basta resistance) was introduced into Agrobacterium strain AGL1, and transformed into conidia of PJ52 according to our previous work (Fan et al., 2019). Positive transformants were confirmed by PCR and subjected to phenotype analysis and disease assay.

To generate transgenic rice plants, Agrobacterium strain EHA105 containing 35S-UvChi1 or Cas9-OsCEBiP/OsCERK1 was introduced into rice accession Q455 or NPB via Agrobacterium-mediated transformation. Positive transgenic lines were confirmed by hygromycin test and PCR.

### Validating secretion of UvChi1

To verify the functionality of UvChi1 signal peptide, plasmids of pSUC2-UvChi1^SP^, pSUC2-Avr1b^SP^, and pSUC2-Mg87^N^ were transformed into yeast strain YTK12 and subjected to yeast secretion assay as performed previously (Fan et al., 2019).

To further confirm whether UvChi1 could be secreted, a multiclonal antibody against UvChi1 in rabbits using the synthetic peptide GRADPSPQGEDLTTSC was raised at Hangzhou Hua’an Biotechnology Co., Ltd, China. *U. virens* PJ52 was cultured in PSB for seven days, and then mycelia and culture supernatant were separated for protein extraction following a previous report (Zhang et al., 2020). The proteins were separated in 10% SDS-PAGE gels and subjected to Western blot using anti-UvChi1 and anti-GAPDH antibodies.

### Chitin binding and chitinase activity assays

The recombinant proteins (GST-UvChi1, MBP-UvChi1, MBP-UvChi1^mcb^, MBP-OsCEBiP, GST, and MBP) were purified from *Escherichia coli* and used for chitin binding assay as described with modifications (Han et al., 2019). Briefly, the recombinant proteins (a final concentration of 0.06 mg ml^−1^) were incubated with chitin beads (a final concentration of 20 μl ml^−1^), shrimp shell chitin (20 mg ml^−1^), cellulose (20 mg ml^−1^), or chitosan (20 mg ml^−1^) in 800 μl ddH_2_O at 4°C. After 4 h, the insoluble pellet fraction was centrifuged (4°C, 12000 rpm, 10 min), and the supernatant was collected. The insoluble pellets were rinsed three times with ddH_2_O. Both the supernatants and the pellets were boiled in 1% SDS for extraction of proteins, which were then separated in 10% SDS-PAGE gels and immunoblotted with anti-GST (Invitrogen) or anti-MBP antibodies (NEB).

Chitinase activity assay was conducted following a previous method with minor modifications (Thompson et al., 2001). Briefly, 20 μl of fluorescent substrate 4-methylumbelliferyl-β-D-N, N ’, N ” -triacetylchitotriose (1 mM in DMSO) was mixed with 150 μl of 200 mM sodium phosphate buffer (pH 6.7), and incubated for 20 min at 37°C. The reaction started by the addition of 30 μl of GST-UvChi1 or GST (2.0 mg ml^−1^ each). After 2 h at 37 ℃, the reaction was stopped by the addition of 50 μl of 3 M NaCO_3_. The fluorescence of released by 4-methylumbelliferone was determined at an excitation wavelength of 390 nm and emission wavelength of 442 nm using spectral scanning multifunctional reader (Thermo Scientific Variskan Flash 4.00.53).

### Nucleic acid extraction, RT-qPCR, and PCR

*U. virens* mycelia were collected from PSA for extraction of genomic DNA and total RNA using CTAB method (Doyle and Doyle, 1990) and TRIzol reagent (Invitrogen), respectively. For quantification of *UvChi1* transcriptional level, *U. virens* RNA was reverse-transcribed using ReverTra Ace PCR RT Kit (TOYOBO); the resultant cDNA was used for qPCR with SYBR Green mix (Qiagen) and gene-specific primers. *UvTub2α* was served as a reference gene. For polymorphism analysis of *UvChi1*, genomic DNA of tested *U. virens* isolates were used to amplify the full-length of *UvChi1* gene. The PCR products were directly sequenced at Sangon Biotech (Chengdu) Co., Ltd, China. Primers used in this study are listed in Supplemental Table S2.

For experiments of chitin co-incubation with the recombinant proteins, fully-expanded rice leaves at 4-leaf stage and developing spikelets at late booting stage were used. Leaf discs or individual spikelets were floated on ddH_2_O overnight at room temperature, and then treated with chitin (30 μg ml^−1^) after 1 h-incubation with the recombinant proteins (GST-UvChi1, GST, MBP-UvChi1, MBP-UvChi1^mcb^, and MBP; 30 μg ml^−1^ for each). Rice samples were then collected for total RNA extraction and RT-qPCR analysis. Note that the recombinant proteins were dialyzed with ddH_2_O before use.

For experiments of leaf co-incubation with the recombinant proteins, leaf discs were floated on ddH_2_O overnight at room temperature, and then co-incubated with the recombinant proteins (MBP-UvChi1, MBP-UvChi1^mcb^, or MBP; 30 μg ml^−1^ for each) for 6 h before chitin treatment. This co-incubation treatment may give enough time for the recombinant proteins to approach the cell surface receptors OsCEBiP and OsCERK1. RT-qPCR analysis was performed using *OsUbi* used as a reference gene. Primers used in this study are listed in Supplemental Table S2.

### Measurement of reactive oxygen species

To determine the burst of ROS in rice or *N. benthamiana* leaves, leaf discs were floated on ddH_2_O overnight. For experiments of chitin co-incubation with the recombinant proteins, chitin (8 μM, hexa-N-acetylchitohexaose) was incubated with the recombinant proteins (GST-UvChi1, GST, MBP-UvChi1, MBP-UvChi1^mcb^, and MBP; 30 μg ml^−1^ for each) for 1h, and then mixed with 20 μM luminol and 10 μg ml^−1^ horseradish peroxidase. The resultants were applied to leaf discs to induce ROS, which was measured in a GloMax 20/20 luminometer (Shi et al., 2018). For experiments of leaf co-incubation with the recombinant proteins, overnight leaf discs were incubated with the recombinant proteins (MBP-UvChi1, MBP-UvChi1^mcb^, or MBP) for 6 h, and then treated with ROS-inducing mixture (8 μM chitin, 20 μM luminol, and 10 μg ml^−1^ horseradish peroxidase).

### GST pull-down assay

The recombinant proteins (GST-OsCEBiP, GST-OsCERK1, MBP-UvChi1, MBP-UvChi1^mcb^, GST-UvChi1, GST-MoChia1, MBP-OsMBL1, MBP-OsTPR1, GST, and MBP) were purified from *E. coli* and used for GST pull-down assays as described previously (Wang et al., 2017). Detection of GST- and MBP-fused proteins was performed with anti-GST (Invitrogen) and anti-MBP antibodies (NEB), respectively.

### Co-immunoprecipitation assay

Agrobacteria GV3101 strains containing indicated constructs were adjusted to a concentration of OD600 = 0.5, and infiltrated into the leaves of *N. benthamiana*. At 36-48 h post infiltration, total proteins of treated leaves were isolated for co-immunoprecipitation assay according to a previous study (Zhou et al., 2015). Anti-FLAG, anti-UvChi1, and anti-GFP antibodies were used for immunoblotting. For Co-IP competition assays, chitin (from shrimp shells, 20 μg ml^−1^) was infiltrated into *N. benthamiana* leaves at 10 min before protein extraction.

### Sequence analysis

The genomic sequences of *OsCEBiP* and *OsCERK1* for each haplotype were retrieved from the database of MBKbase-rice (http://mbkbase.org/rice). The genomic sequence of *UvChi1* was amplified from different *U. virens* isolates (Supplemental Table S1) with gene-specific primers (Supplemental Table S2) and identified by sequencing. Corresponding protein sequences were deduced by using online software (https://web.expasy.org/translate). Sequence alignment was conducted with the MultAlin software (Corpet, 1988).

## Supplemental Data

The following materials are available in the online version of this article.

**Supplemental Figure S1.** *UvChi1* encodes a fungal chitinase.

**Supplemental Figure S2.** Generation of *UvChi1* knockout mutants and pathogenicity test.

**Supplemental Figure S3.** Ectopic expression of *UvChi1* promotes *Ustilaginoidea virens* infection in rice.

**Supplemental Figure S4.** UvChi1 binds to chitin and blocks chitin perception in rice.

**Supplemental Figure S5.** UvChi1 does not interact with OsMBL1 and OsTPR1.

**Supplemental Figure S6.** *OsCEBiP*, *OsCERK1*, *OsMBL1*, and *OsTPR1* are highly expressed in rice spikelets.

**Supplemental Figure S7.** Generation of UvChi1 protein mutated at chitin-binding sites.

**Supplemental Figure S8.** Generation of *oscebip*/*oscerk1* double mutants.

**Supplemental Figure S9.** Protein sequence alignment of OsCEBiP^58-356^.

**Supplemental Figure S10.** UvChi1 interferes with the oligomerizations of OsCEBiP^T2^ and OsCERK1^T2^.

**Supplemental Figure S11.** A proposed model of UvChi1 virulence mechanisms.

**Supplemental Table S1.** *Ustilaginoidea virens* isolates used for DNA polymorphism analysis.

**Supplemental Table S2.** Primers used in this study.

## ACKNOWLEDGEMENTS

We thank Dr. Yongfeng Liu for pSK1044 plasmid, Zhengguang Zhang for pCas9-tRP-gRNA vector, and Pengfei Qi for pRF-HU2 vector. We thank Drs. Dongwei Hu, Chaoxi Luo, Peizhou Xu, and Chaoqun Zang for providing *U. virens* isolates used for polymorphism analysis.

