## Supplemental Fig for "Insights into the susceptibility of rice to a floral disease"

### SUPPLEMENTAL FIGURES

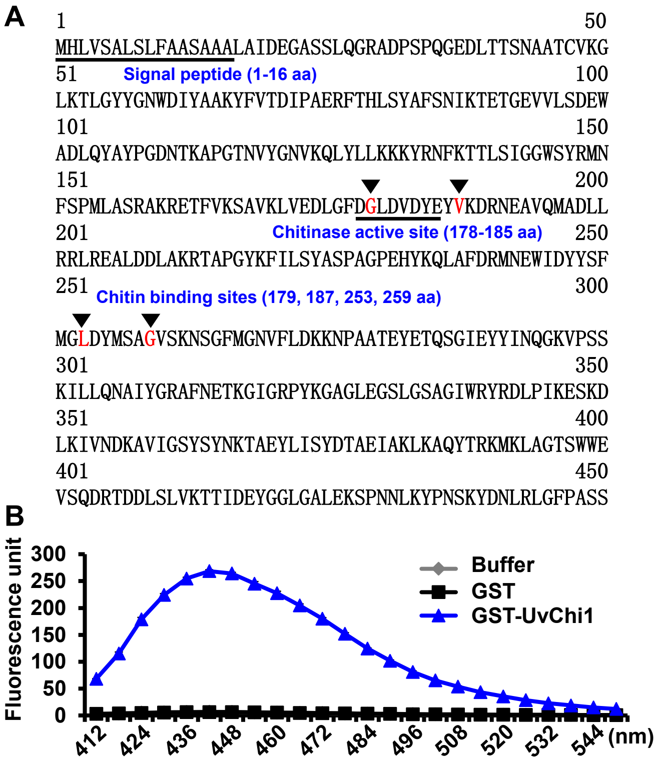


**Supplemental Figure S1.** *UvChi1* encodes a fungal chitinase. **A,** Amino acid sequence of UvChi1. Signal peptide and chitinase active site are underlined. Chitin binding sites are indicated in red letters and marked by inverted black triangles. **B,** Chitinase activity assay of UvChi1. The recombinant GST-UvChi1 was purified from *Escherichia coli* and incubated with 4-Methylumbelliferyl-ß-D-N,N'N'-triacetyl- chitotriose (MUC3). GST-UvChi1 could degrade nonfluorescent MUC3 into fluorescent 4-muthylumbelliferone, while the GST protein or buffer could not. Data are represented as mean ± SD of three biological replicates.

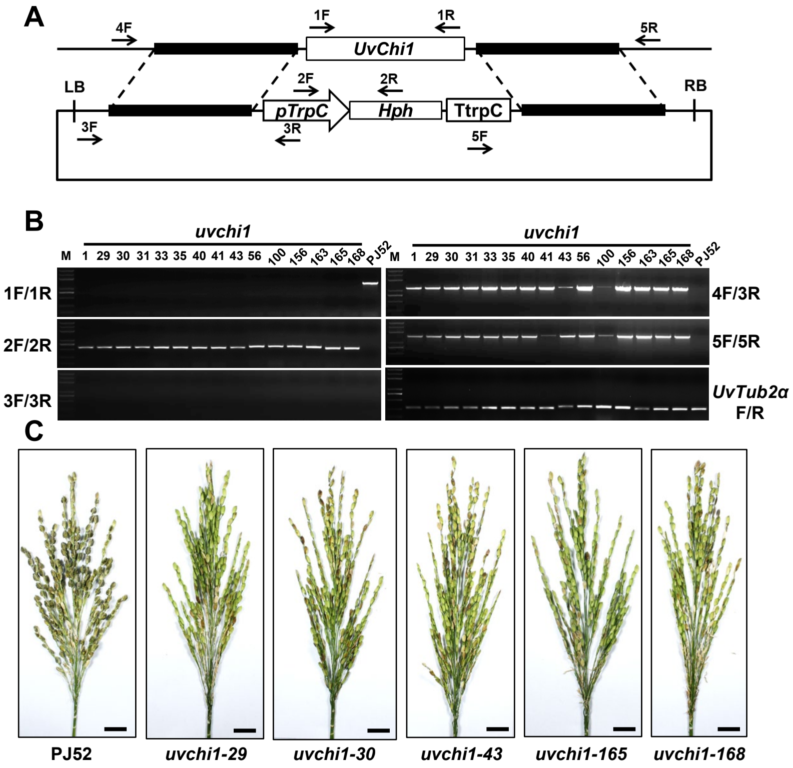


**Supplemental Figure S2.** Generation of *UvChi1* knockout mutants and pathogenicity test. **A,** Schematic diagram of replacing *UvChi1* with TrpC::hph resistance cassette for generation of *uvchi1* mutants. Primers for confirmation of *uvchi1* mutants were indicated as arrows. **B,** Electrophoresis analysis of PCR products amplified by multiple primer pairs in **(A)**. M, Trans2K Plus DNA Marker. **C,** False smut disease assay. Indicated strains were inoculated into a susceptible rice accession Q455 ($n$ > 10). Disease phenotype was recorded at four week post inoculation. Note that no false smut balls were developed from *uvchi1* mutants-inoculated panicles. Size bar, 2 cm.

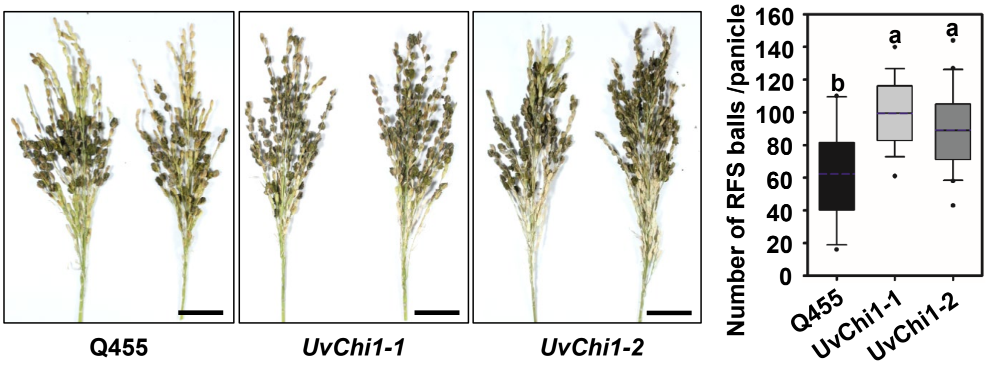


**Supplemental Figure S3.** Ectopic expression of *UvChi1* promotes *Ustilaginoidea virens* infection in rice. Disease assay of transgenic rice ectopically expressing *UvChi1*. *U. virens* PJ52 was inoculated into Q455 and transgenic lines *UvChi1-1* and *UvChi1-2*. Disease phenotype was recorded at four week post inoculation, and the number of false smut balls was statistically analyzed. Data are box-plotted ($n > 30$). Different letters above data box indicate significant differences determined by one-way ANOVA with post hoc Tukey HSD analysis ($P < 0.05$). Size bar, 2 cm.

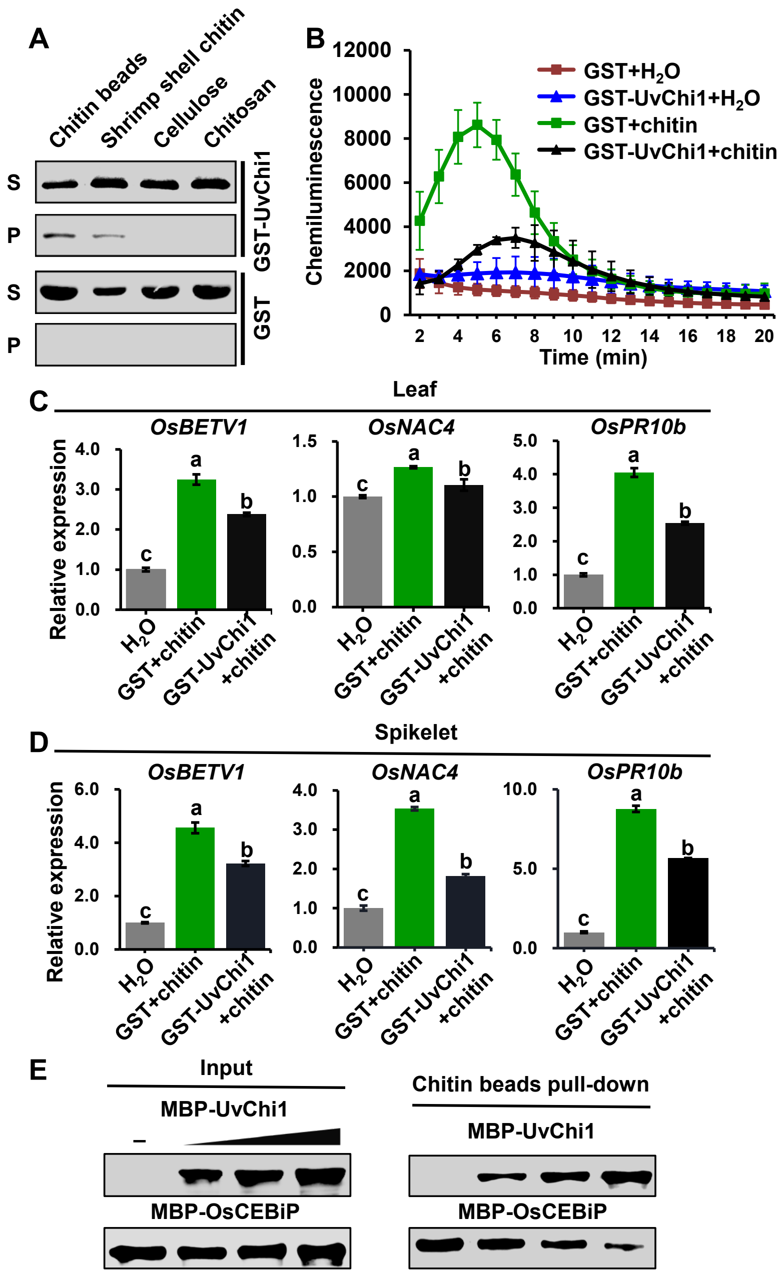


**Supplemental Figure S4.** UvChi1 binds to chitin and blocks chitin perception in rice. **A,** Chitin binding assay of UvChi1. The recombinant protein GST-UvChi1 was incubated with insoluble chitin beads, shrimp shell chitin, cellulose, and chitosan. GST protein was used as the control. After precipitation, pellets (P) and supernatants (S) were detected with anti-GST antibody. **B,** Chitin-induced ROS assay of rice leaves treated with GST-UvChi1. Chitin was incubated with GST-UvChi1 or GST for 1 h, and then the resultant was used for inducing ROS of rice leaf discs. Data are represented as mean ± SD of three biological replicates. Experiments were repeated twice with similar results. **C-D,** Chitin-induced expression of defense-related genes in rice leaf **(C)** or spikelet **(D)**. Chitin was incubated with GST-UvChi1 or GST for 1 h, and then the resultant was used to treat rice leaves (for 12 h) or spikelets (for 6 h). Relative expression of indicated genes was determined using *OsUbi* as the reference gene. Data are represented as mean ± SD of three replicates. Different letters above data box indicate significant differences determined by one-way ANOVA with post hoc Tukey HSD analysis ($P < 0.05$). Experiments were repeated twice with similar results. **E,** UvChi1 competes with OsCEBiP for chitin binding. Increasing amounts of MBP-UvChi1 was mixed with MBP-OsCEBiP, and the resultants were incubated with chitin beads. MBP-UvChi1 or MBP-OsCEBiP pull-down by chitin beads was detected with anti-MBP antibody.

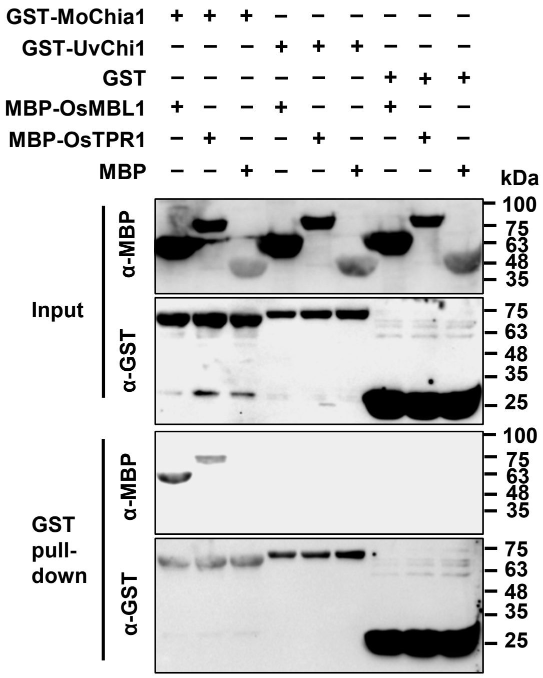


**Supplemental Figure S5.** UvChi1 does not interact with OsMBL1 and OsTPR1. The recombinant proteins GST-MoChia1, GST-UvChi1, MBP-OsMBL1, and MBP-OsTPR1 were purified from *Escherichia coli*, and subjected to GST pull-down assays. GST and MBP tag proteins were used as negative controls. GST-MoChia1 that interacts with MBP-OsMBL1 and MBP-OsTPR1 was served as the positive control.

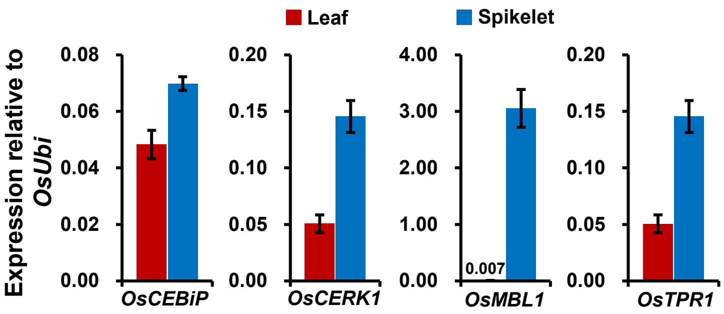


**Supplemental Figure S6.** *OsCEBiP*, *OsCERK1*, *OsMBL1*, and *OsTPR1* are highly expressed in rice spikelets. Rice leaf and spikelet samples were collected for RT-qPCR analysis of *OsCEBiP*, *OsCERK1*, *OsMBL1*, and *OsTPR1*. Relative expression levels of indicated genes were calculated relative to that of *OsUbi*. Data are represented as mean ± SD of three biological replicates.

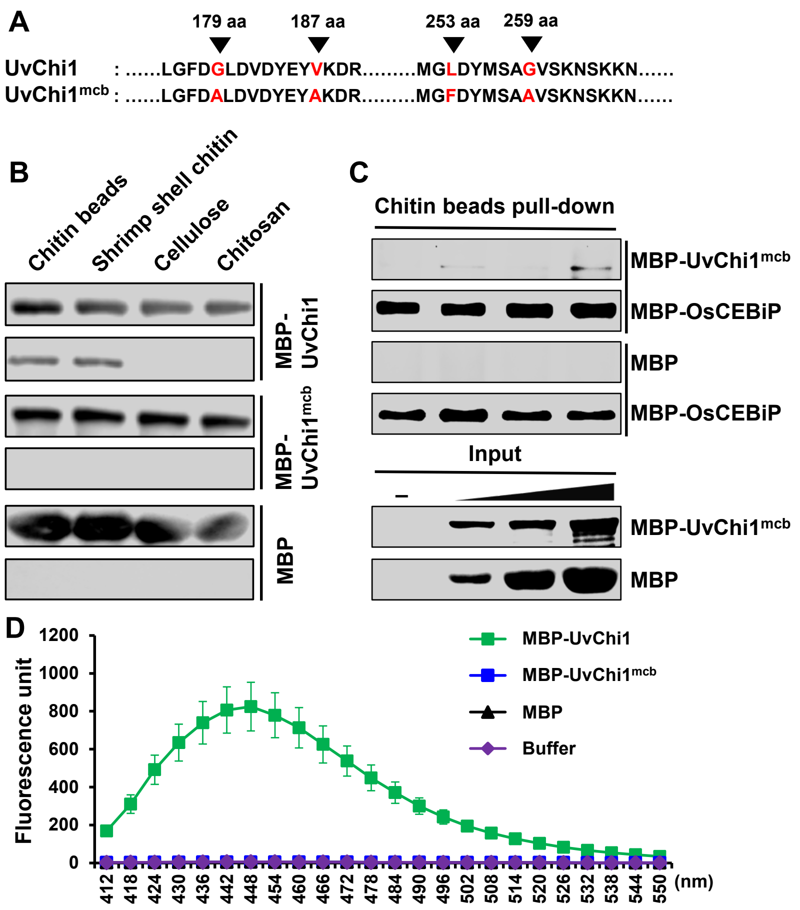


**Supplemental Figure S7.** Generation of UvChi1 protein mutated at chitin-binding sites. **A,** Chitin binding sites of UvChi1 were mutated to generate UvChi1$^{mcb}$ (G179A, V187A, L253F, and G259A). **B,** Chitin binding assay of UvChi1 and UvChi1$^{mcb}$. The recombinant proteins MBP-UvChi1 and MBP-UvChi1$^{mcb}$ were respectively incubated with insoluble chitin beads, shrimp shell chitin, cellulose, and chitosan. MBP tag protein was used as the control. After precipitation, pellets (P) and supernatants (S) were detected by Western blot with anti-MBP antibody. **C,** UvChi1$^{mcb}$ does not interfere with OsCEBiP binding to chitin. Increasing amounts of MBP or MBP-UvChi1$^{mcb}$ were mixed with MBP-OsCEBiP, and the resultants were incubated with chitin beads. MBP, MBP-UvChi1$^{mcb}$, and MBP-OsCEBiP was detected with anti-MBP antibody after pull-down by chitin beads. **D,** Chitinase activity assay of UvChi1$^{mcb}$. The assay was conducted as described in Supplemental Figure S1B. Data are represented as mean ± SD of three replicates.

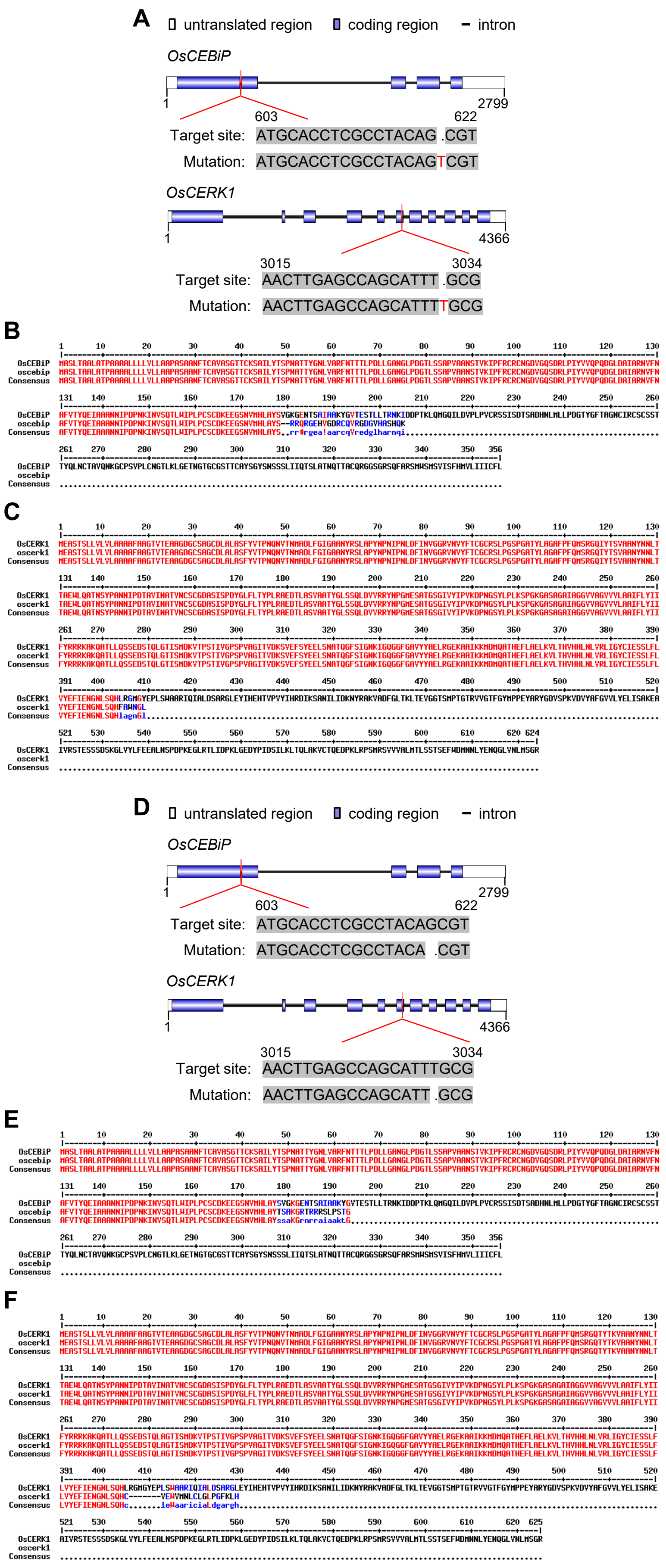
A
untranslated region
coding region
intron
OsCEBiP
1
2799
603
622
Target site: ATGCACCTCGCCTACAG .CGT
Mutation: ATGCACCTCGCCTACAGTCGT
OsCERK1
1
4366
3015
3034
Target site: AACTTGAGCCAGCATTT .GCG
Mutation: AACTTGAGCCAGCATTTTGCG
B
OsCEBiP
oscebip
Consensus
C
OsCERK1
oscerk1
Consensus
D
untranslated region
coding region
intron
OsCEBiP
1
2799
603
622
Target site: ATGCACCTCGCCTACAGCGT
Mutation: ATGCACCTCGCCTACA .CGT
OsCERK1
1
4366
3015
3034
Target site: AACTTGAGCCAGCATTTGCG
Mutation: AACTTGAGCCAGCATT .GCG
E
OsCEBiP
oscebip
Consensus
F
OsCERK1
oscerk1
Consensus

**Supplemental Figure S8.** Generation of *oscebip*/*oscerk1* double mutants. **A and D,** CRISPR/Cas9 approach was applied to simultaneously knockout *OsCEBiP* and *OsCERK1* in Nipponbare (NPB; **A**) and Q455 (**D**), by designing one target site on each gene. The sequences and positions of target sites are presented. **B and E,** Protein sequence alignment of OsCEBiP in NPB (b), Q455 (e) and *oscebip*/*oscerk1* mutants. **C and F,** Protein sequence alignment of OsCERK1 in NPB **(C)**, Q455 **(F)** and *oscebip*/*oscerk1* mutants. OsCEBiP and OsCERK1 proteins were both truncated in *oscebip*/*oscerk1*.

```
P1   --LVARFNTTTLPDLLGANGLPDGTLSSAPVAANSTVKIPFRCRCNGDVGQ
P2   --LVARFNTTTLPDLLGANGLPDGTLSSAPVAANSTVKIPFRCRCNGDVGQ
P3   --SSPASTPPPSPTSSAPTASPTARFPPPPSPPIPPSKSPSAAAATATSAS
P4   PRRPLQHHHPPRPPRRQRPPRRHAFLRPRRRQFHRQNPLPLPLQRRRRPVG

P1   SDRLPIYVVQPQDGLDAIARNVFNAFVTYQEIAANNIPDPNKINVSQTLW
P2   SDRLPIYVVQPQDGLDAIARNVFNAFVTYQEIAPRTTSPTPTR-------
P3   RTASPSTSCSRRTGSTPSRATCSTPSSPTRRSPPRTTSPTPTR-------
P4   PPPHLRRAAAGRARRHRAQRVQRLRHLPGDR-QPRTTSPTPTR-------

P1   IPLPCSCDKEEGSNVMHLAYSVGKGENTSAIAAKYGVTESTLLTRNKIDDP
P2   --------------------------------------------------
P3   --------------------------------------------------
P4   --------------------------------------------------

P1   TKLQMGQILDVPLPVCRSSISDTSADHNLMLLPDGTYGFTAGNCIRCSCSS
P2   --------------------------------------------------
P3   --------------------------------------------------
P4   --------------------------------------------------

P1   TTYQLNCTAVQNKGCPSVPLCNGTLKLGETNGTGCGSTTCAYSGYSNSSSL
P2   --------------------------------------------------
P3   --------------------------------------------------
P4   --------------------------------------------------

P1   IIQTSLATNQTTACQRGGSGRSQFARSMWSMSVISFHMVLIIICFL
P2   ----------------------------------------------
P3   ----------------------------------------------
P4   ----------------------------------------------
```

**Supplemental Figure S9.** Protein sequence alignment of OsCEBiP$^{58\text{-}356}$. As depicted in Figure 5A, polymorphic types of OsCEBiP C-terminus (at position of 58-356 aa) in over 5000 rice accessions are classified into four types, including P1, P2, P3, and P4. OsCEBiP$^{58\text{-}356}$ sequence alignment for P1-P4 was presented.

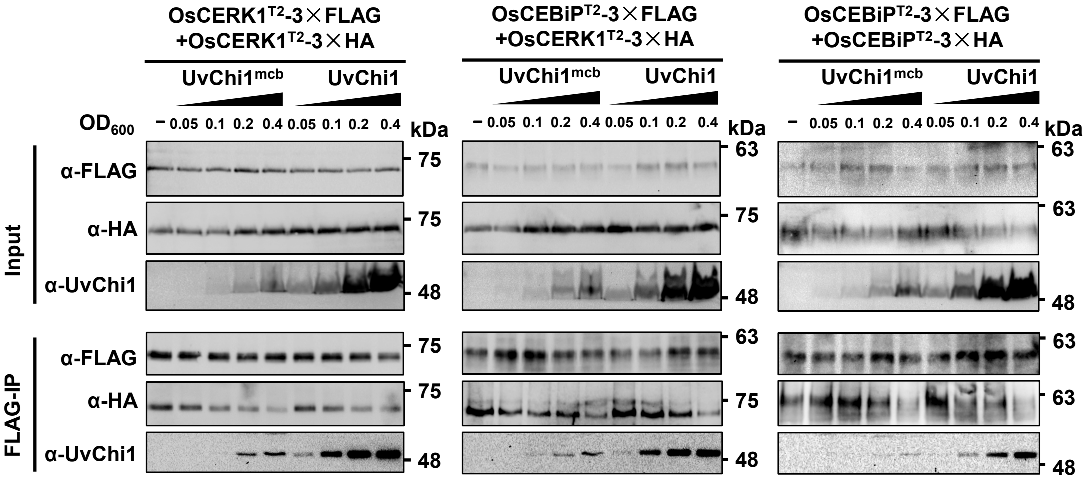


**Supplemental Figure S10.** UvChi1 interferes with the oligomerizations of OsCEBiP$^{T2}$ and OsCERK1$^{T2}$. *In vivo* Co-IP competition assays were conducted as described in Figure 4A.

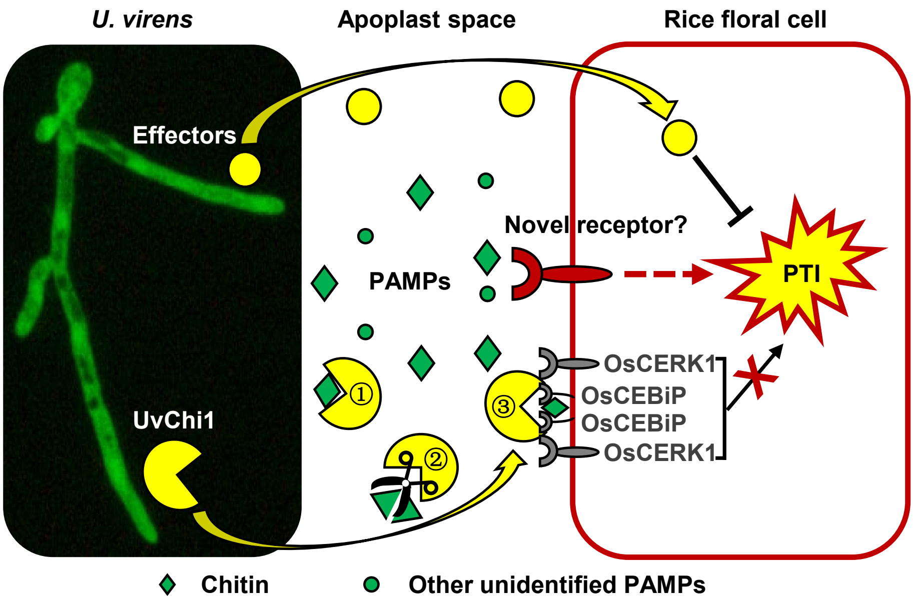


**Supplemental Figure S11.** A proposed model of UvChi1 virulence mechanisms. *U. virens* can secrete a series of effector proteins during infection of rice flower. In addition to immunosuppressive effectors that may be translocated into rice cells (Fan *et al.*, 2019; Fang *et al.*, 2019; Zhang *et al.*, 2020), *U. virens* also secretes apoplastic effectors such as UvChi1 that interferes with chitin perception and signaling in rice. UvChi1 may binds to (①) and/or degrade (②) chitin to sequester chitin-triggered rice immunity. Also, UvChi1 can interact with chitin receptor OsCEBiP and co-receptor OsCERK1 to interfere with their interactions and downstream immune responses (③). Accordingly, rice may employ other chitin sensors to compete with UvChi1 for chitin binding, or novel receptors to perceive *U. virens*, leading to regain of rice immunity against *U. virens*.

#### SUPPLEMENTAL TABLES (*provided as a separate Excel file*)

**Supplemental Table S1.** *Ustilaginoidea virens* isolates used for DNA polymorphism analysis.

**Supplemental Table S2.** Primers used in this study.
